# Characterization of Composite Agarose-Collagen Hydrogels for Chondrocyte Culture

**DOI:** 10.1101/2024.03.02.583023

**Authors:** Clarisse Zigan, Claudia Benito Alston, Aritra Chatterjee, Luis Solorio, Deva D. Chan

**Author notes:** Corresponding Author: Deva Chan.

## Abstract

To elucidate the mechanisms of cellular mechanotransduction, it is necessary to employ biomaterials that effectively merge biofunctionality with appropriate mechanical characteristics. Agarose and collagen separately are common biopolymers used in cartilage mechanobiology and mechanotransduction studies but lack features that make them ideal for functional engineered cartilage. In this study, agarose (8% w/v and 4% w/v) is blended with collagen type I (4mg/mL) to create composites. We hypothesized that a higher stiffness, composite hydrogel would promote native cartilage-like conditions. To address these questions, acellular and cell-laden studies were completed to assess rheologic and compressive properties, contraction, and structural homogeneity in addition to matrix mechanics, cell proliferation, and glycosaminoglycan production. Over 21 days in culture, cellular 4% agarose – 2mg/mL collagen I hydrogels displayed good structural and bulk mechanical properties, cell proliferation, and continual glycosaminoglycan production, indicating promise towards the development of an effective hydrogel for chondrocyte mechanotransduction and mechanobiology studies.

**Graphical Abstract:** 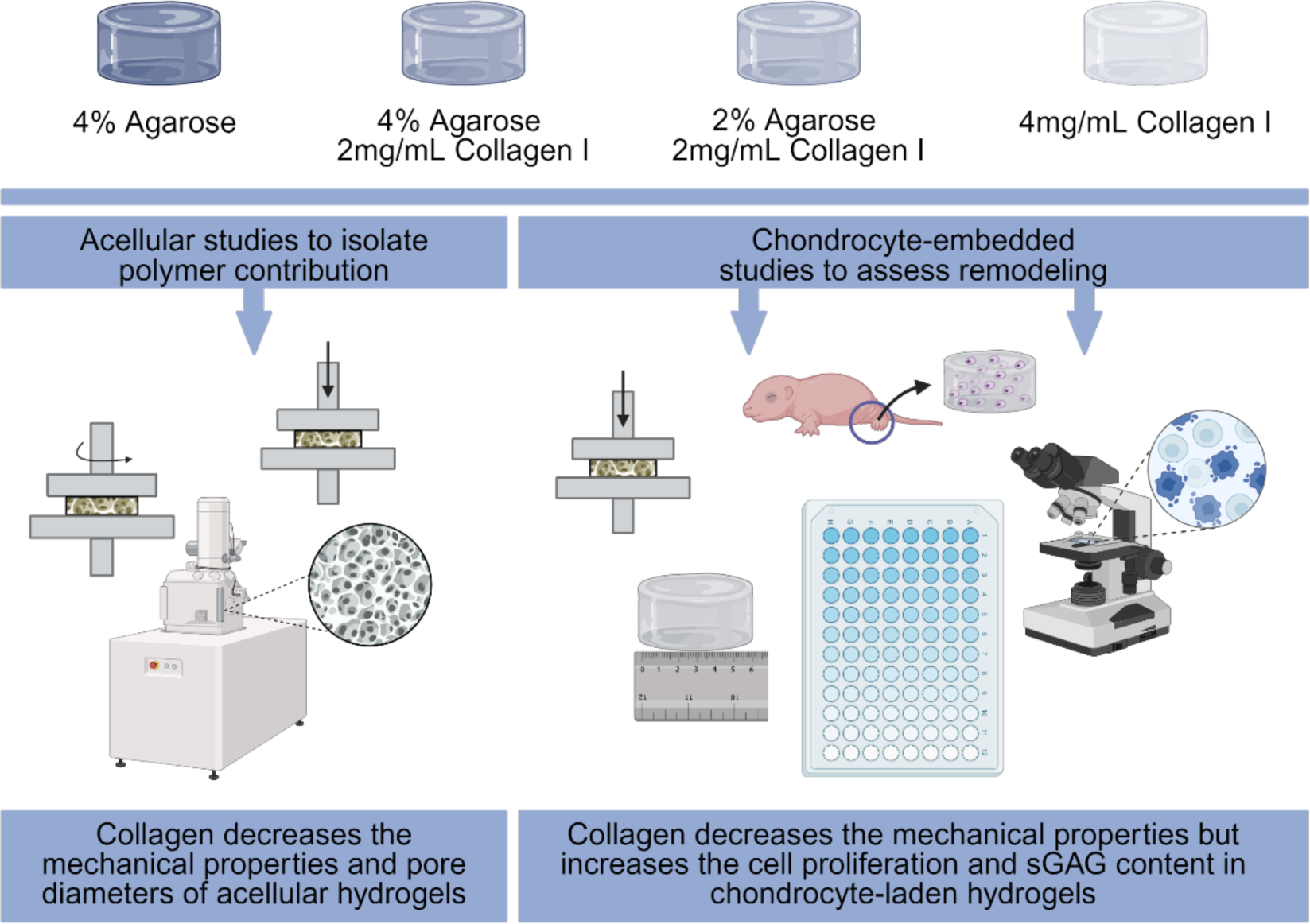

## INTRODUCTION

Cartilage mechanobiology and tissue engineering fields have both evolved using strategies employed to repair or regenerate damaged cartilage using a combination of biomaterials, cells, and stimulation. It is well accepted that applying mechanical forces to engineered cartilage constructs can help mimic the natural environment and stimulate chondrocytes to produce extracellular matrix (ECM) components and maintain tissue integrity to enhance the formation of functional cartilage tissue ^1–3^. However, the underlying mechanisms that are employed during these processes are poorly understood ^4,5^. Many studies focus primarily on the effects of loading conditions applied to cartilage constructs ^6–10^. However, another key factor to consider is construct composition, since administered mechanical forces can be influenced by exogenous physical and chemical cues of the 3-dimensional matrix that envelops the cell ^11^. Cartilage-like constructs are often formed of polymeric hydrogels ^12,13^.

Agarose is a common natural polymer used for cartilage mechanobiology studies due to its easily tunable mechanical stiffness. However, prior chondrocyte biology studies use low (2-3% v/v) concentration agarose-based hydrogels ^14–18^, which do not achieve cartilage-like mechanical properties as their corresponding moduli range from 13 kPa to 18 kPa. Zignego, *et. al.,* however, found that higher concentration 4.5% w/v agarose hydrogels achieve stiffness levels within ranges found in the native pericellular matrix (PCM; 20-200 kPa ^19,20^) while still maintaining chondrocyte viability, highlighting potential of high stiffness microenvironments to better match cartilage properties for subsequent mechanotransduction studies ^21^. However, without modification, agarose lacks integrin binding motifs that are critical in the mediation of cell-ECM interactions.

Collagen I is another natural polymer common in chondrocyte studies ^22–24^. Unlike agarose, collagen offers ample binding motifs ^25^, to promote cell adhesion and migration. Collagen hydrogels have a large water content leading to flexibility similar to natural tissue, exhibit biocompatibility and biodegradation abilities, and can be mechanical tuned through physical or chemical cross-linking ^12^. While type II collagen is the most abundant extracellular molecule in cartilage, providing foundation for the use of collagen hydrogels, *in vitro* studies generally use type I collagen hydrogels since they demonstrate superior mechanical qualities. However, type I collagen hydrogels are still soft compared to cartilage and often do not maintain their 3-dimensional structure for extended studies ^22^. For example, 1-3 mg/mL concentrations, a typical range found in chondrocyte-embedded collagen hydrogel studies, result in elastic moduli of 5-22 pascals ^22,26,27^.

Composite hydrogels combine the favorable properties of each individual polymer, enabling tunable mechanical and structural properties and leading to more ECM-like interactions that promote cues for proliferation, differentiation, and matrix production. Few groups have studied the interactions between agarose and collagen biomaterials and their influence on cells ^28–33^. Cambria, *et. al.,* assessed the impact of low concentration agarose blended with collagen on nucleus pulposus cells, finding composite hydrogels to outperform agarose-only hydrogels in terms of cell adhesion and proliferation, likely attributable to the binding motifs that exist within the collagen component ^28^. Quarta, *et. al.,* cultured breast cancer cell lines in agarose-collagen hydrogels to assess the mechanical properties and potential cytotoxicity as agarose content was increased, of which at the low concentrations assessed, while there was a change in mechanical behavior, no cytotoxicity was observed ^29^. Ulrich, *et. al.,* used another cancer cell line to assess how an increase in hydrogel stiffness would alter cell motility and therefore the potential influence of agarose on collagen deformation and remodeling ^30^. While the group’s work with mass spectrometry and scanning electron microscopy fail to indicate agarose induces collagen ligand alterations, microscopy images do suggest agarose inhibits cell-directed assembly of large collagen bundles, also influencing cell spreading and motility ^30^. While these studies demonstrate applications of agarose-collagen hydrogels and their compositional influence on mechanics, cell viability, and remodeling, study of the response of chondrocytes to this environment are critical to the application of such hydrogels to the study of cartilage mechanobiology.

In this study, we aimed to develop an agarose-collagen composite hydrogel that combines the mechanical properties of agarose with the biofunctionality of collagen to mimic native articular cartilage tissue and enable future chondrocyte mechanobiology studies. Comparing against agarose and collagen only hydrogels, we evaluated agarose-collagen composite hydrogel mechanical properties and structural homogeneity alongside their effect on chondrocyte viability, proliferation, and sulfated glycosaminoglycan (sGAG) production.

## MATERIALS & METHODS

Experiments conducted in this study were divided into two main categories (1) acellular and (2) cell laden. All hydrogel formulations were otherwise prepared in the same fashion. The aim of acellular experiments was to elucidate the innate structure-function relationships between the hydrogel formulation and the resulting bulk mechanics and structural homogeneity. Cell-laden hydrogel experiments were then used to assess chondrocyte morphology, proliferation rates, and sGAG production.

### Hydrogel Preparation

To prepare agarose hydrogels, low-gelling temperature type VII-A agarose (A0701, Sigma Aldrich) was dissolved in 1× phosphate buffered saline (PBS) and autoclaved (chamber temperature of 120°C and sterilization time set to 15 minutes). For cell-based experiments, the solution was cooled to 40°C prior to adding chondrocytes and casting in the mold.

Collagen I hydrogels were prepared by chilling and mixing rat tail collagen I (RatCol®; 5153, Advanced BioMatrix) with its associated neutralizing solution in accordance with the manufacturer protocol to obtain a 4mg/mL solution.

Two composite hydrogel formulations were studied. The first composite consisted of 4% w/v agarose with 2 mg/mL collagen I. The second composite consisted of a 1:1 ratio of the aforementioned agarose-only and collagen-only formulations, leading to 2% w/v agarose with 2 mg/mL collagen I concentration. Composite hydrogels were manually mixed via pipetting while maintained in a ∼40°C water bath to prevent collagen denaturation and premature agarose gelation.

Positive displacement pipette tips were used to aliquot 150-μL volumes of each hydrogel solution into custom (3-mm tall, 6-mm inner diameter) PDMS based molds (00-30, Ecoflex).

### Cell Encapsulation

To obtain primary chondrocytes, seven C57BL/6J mice (5 days old) were humanely euthanized under institutional approval (IACUC protocol 2104002138) to isolate neonatal cartilage of the proximal and distal femur and the proximal tibia under sterile conditions. Tissues were digested in 3 mg/mL type II collagenase (17101-015, Gibco) reconstituted in culture media for one hour at 37°C followed by digestion in 0.5 mg/mL type II collagenase overnight at 37°C. Tissue was agitated to dissociate residual tissue pieces, and the entire solution was filtered through a 40-μm cell strainer into a 50-mL conical tube. Sterile 1× PBS was added until a 40-mL total volume was reached, at which point the solution was centrifuged at 1000 rpm for 10 minutes. The supernatant was aspirated, and the cell pellet resuspended in 30 mL of sterile 1× PBS to be re-centrifuged at 1000 rpm for 10 minutes. The supernatant was again aspirated, and the pellet resuspended in complete growth media and counted.

Chondrocytes were resuspended into hydrogels at concentrations of 1 × 10^6^ cells/mL prior to full hydrogel gelation. Hydrogels with and without cells were allowed to solidify at room temperature for 10 minutes, followed by the addition of 0.5-mL complete culture media [phenol-free 4.5 g/L glucose Dulbecco’s modified Eagle medium (DMEM; 31053-036, Gibco) supplemented with L-glutamate (AAJ6057322, Gibco) and sodium pyruvate (11360-070, Gibco)) completed with 10% fetal bovine serum (FBS; 12676029, Corning) and 1% penicillin-streptomycin (15140-122, Gibco)]. Hydrogels were cultured in a 37°C, 5% CO2 incubator in 48-well plates for up to 21 days. Culture medium was changed every 2-3 days.

### Rheological Characterization of Acellular Hydrogels

Following recommendations for rheological characterization of hydrogels ^34^, small amplitude oscillatory shear tests were performed (AR-G2 Rheometer, TA Instruments) using a 20-mm diameter parallel plate geometry and hydration chamber to mitigate evaporation. 150-µL acellular hydrogel solution was dispensed between plates. After shear rheometric parameters were set (Table 1), the plate was rapidly heated or cooled to begin gelation. Storage (*G*′) and loss (*G*^!!^) moduli were defined as the average of all values recorded per step.

**Table 1.**
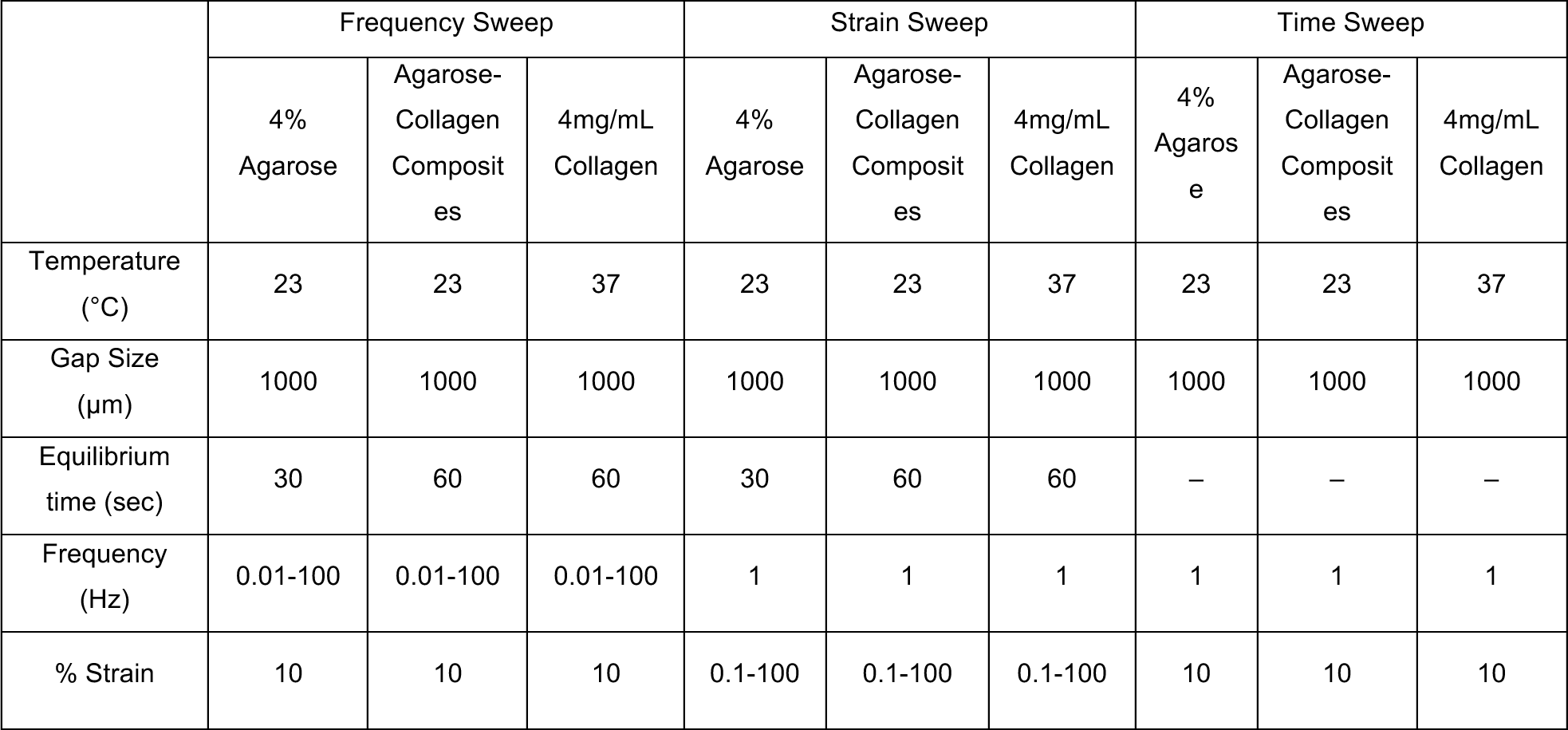
Parameters for shear rheometric testing.

|  | Frequency Sweep |  |  | Strain Sweep |  |  | Time Sweep |  |  |
| --- | --- | --- | --- | --- | --- | --- | --- | --- | --- |
|  | 4% Agarose | Agarose-Collagen Composites | 4mg/mL Collagen | 4% Agarose | Agarose-Collagen Composites | 4mg/mL Collagen | 4% Agarose | Agarose-Collagen Composites | 4mg/mL Collagen |
| Temperature (°C) | 23 | 23 | 37 | 23 | 23 | 37 | 23 | 23 | 37 |
| Gap Size (μm) | 1000 | 1000 | 1000 | 1000 | 1000 | 1000 | 1000 | 1000 | 1000 |
| Equilibrium time (sec) | 30 | 60 | 60 | 30 | 60 | 60 | – | – | – |
| Frequency (Hz) | 0.01-100 | 0.01-100 | 0.01-100 | 1 | 1 | 1 | 1 | 1 | 1 |
| % Strain | 10 | 10 | 10 | 0.1-100 | 0.1-100 | 0.1-100 | 10 | 10 | 10 |

A frequency sweep between 0.01- and 100-Hz was used to evaluate the crosslinking behavior of the hydrogels. Strain sweeps from 0.1 to 100% strain were used to assess the linear viscoelastic region (LVR) limits on fully formed gels. A time sweep was used to confirm consistent mechanical behavior over time and that no unexpected increase in loss or storage modulus would occur due to innate material properties during culture. Each sample (n=3) was only used for one sweep.

### Unconfined Compression of Hydrogels

The effect of agarose and collagen concentrations on equilibrium moduli were evaluated by uniaxial unconfined compression tests. Bulk mechanical changes were assessed in both acellular (n=6; after 24 hours gelation) and cell-laden hydrogels (n=4; throughout 21-days of culture) using the same protocol. First, the diameter and heights per each individual sample were measured with a digital caliper prior to testing on a universal testing machine (ElectroForce 5500, TA Instruments) equipped with a 20-lbf load cell then maintained in a 1× PBS bath during tests. Stress-relaxation was then performed to 10% strain and held (acellular = 1200 seconds; cell-laden = 600 seconds) to evaluate the compressive modulus and time-dependent behavior (10% strain, 1% strain/sec). The load vs time data obtained from these tests were converted to stress vs time data by applying sample geometry information. These data were then used to estimate the viscoelastic properties of the hydrogels using a nonlinear Prony Series model ^35^:

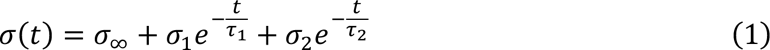

where *σ*_*i*_ and *τ*_*i*_ are stress parameters and relaxation time constants, respectively. From these outputs, the equilibrium modulus (*E*_∞_ = *σ*_∞_/ε) and instantaneous modulus (*E*_0_ = (*σ*_∞_ + *σ*_1_ + *σ*_2_)/ε) could be calculated by normalizing the experienced stresses to the applied strain ^36^. The model was fitted to experimental data using a non-linear least squares method in MATLAB (R2022a, MathWorks).

### Electron Microscopy of Acellular Hydrogels

Texture and homogeneity of hydrogels (n=3) were analyzed by field emission scanning electron microscopy (FE-SEM; SEM). Samples were flash frozen in liquid nitrogen, fractured with a frozen razor blade, and stored at -80 °C. Hydrogels were lyophilized (VirTis, SP Scientific) for 48 hours at -20°C followed by 10 hours at 20°C. Samples were sputter coated with 24-nm of Au-Pd (SPI Supplies) then imaged using a cold field emission high resolution scanning electron microscope (S- 4800, Hitachi) at an operating voltage of 10-kV. SEM images at 100× magnification were used to quantify porosity using an adapted open-source MATLAB script ^37^ while 10,000× SEM images were used to quantify collagen fiber diameters using an adapted open-source Fiber Diameter Distribution v1.0.3 script ^38,39^. In short, to quantify porosities, the image segmentation code segmented images using adaptive thresholding, relying on the mean intensity of a local neighborhood rather than global histogram-based thresholding (i.e., Otsu’s method). An iterative refinement process was applied through erosion and dilation to enhance the segmentation of the region. However, due to the distinct variations between the collagen fibers and background in the 10,000× images, mean and Gaussian filters were applied. Subsequently, column and row sweeping operations were carried out. Additionally, 2000X SEM images were used for qualitative comparison.

### Cell Growth

As a surrogate to evaluate cell viability and proliferation, a resazurin assay was used at the initial timepoint (immediately following cell-laden hydrogel gelation) and intermittently over 21 days of culture. Hydrogels (n=3) were incubated for 4 hours in a solution of FBS-free DMEM supplemented with 10% Resazurin dye (AR002, R&D Systems) at 37°C, 5% CO2. Fluorescence was quantified using an excitation of 530/15 nm and emission of 590/15 nm on a fluorescent plate reader (BioTek Cytation 5, Agilent Technologies).

### Nuclear Morphology

Hydrogels were fixed in 4% paraformaldehyde overnight and stored at 4°C in 1× PBS until ready for immunofluorescent imaging. Hydrogel sample preparation and imaging was completed within 1-week of initial storage. Samples were embedded within a 3% agarose (BP160-100, Fisher Bioreagents) mixture and sectioned in a vibratome (VT 1200, Leica) into 300-µm thick sections. Before microscopic examination, sections were stained with 4,6-diamidino-2-phenylindole (DAPI; H-1200- 10, Vector Laboratories), to detect nuclear morphology and 2-dimensional spatial positioning using an upright confocal (Confocal 800, Zeiss LSM). Images (n=4) were taken of each hydrogel type at random locations within the sample and FIJI were used to analyze morphological properties of the nuclei, including aspect ratios and areas.

### Extracellular Matrix Production

The sGAG content in cell-laden hydrogel constructs was measured to compare ECM remodeling with respect to hydrogel formulations. After 3, 7, 14, or 21 days of culture, hydrogels (n=3) were weighed, flash frozen, lyophilized overnight, and reweighed. Hydrogels were digested in 1 mL of 50 µg/mL Proteinase K (P6556, Sigma Aldrich) in 50 mM Tris, 1 mM CaCl2, pH = 8 for 16 hours at 56°C followed by 30 minutes at 90°C and an additional digest of 4 units of beta-agarase (M0392S, New England BioLabs) for 1 hour at 65°C to ensure full agarose breakdown. Supernatants were collected for dimethylmethylene blue (DMMB) assays, and chondroitin sulfate from bovine trachea (C9819, Sigma Aldrich) was used as a standard. DMMB solution was added (200 µL/well) and absorbance was measured at 540 nm and 590 nm using a plate reader (BioTek Cytation 5, Agilent Technologies). For all samples, sGAG quantity was normalized against hydrogel dry weight.

### Statistical Analysis

Results are reported as mean ± standard error (SE) for tests using at least three replicates. Two-sided, unpaired t-tests were used in rheometry analyses to detect differences in hydrogel plateau or transition points, in SEM imaging to detect pore diameter differences, and in fluorescence imaging to detect nuclear morphology differences. One-way ANOVA with post-hoc Bonferroni corrections were used to detect temporal influences in addition to hydrogel formulation influences during compression tests and sGAG tests. *p-*values less than 0.05 were considered statistically significant. Statistical analyses were performed with MATLAB (R2022a, MathWorks).

## RESULTS

### Agarose Dominates the Mechanical Characteristics of Composite Hydrogels

For all hydrogels, the presence of a crossover point indicated reversible crosslinking (Figure 1). 4 mg/mL collagen hydrogels reach this crossover at the lowest frequency (6.07±1.48 Hz), followed by the two intermediate composites (4% agarose – 2 mg/mL collagen: 10.18±1.34 Hz and 2% agarose – 2 mg/mL collagen: 8.37±1.74 Hz), then 4% agarose hydrogels (12.81±1.69 Hz). At frequencies above 10 Hz (10 – 100 Hz), variations in sample responses increase rapidly. The relationship between the storage and loss moduli trends indicated that at low frequencies, the samples exhibit gel-like behavior whereas at higher frequencies, all begin to display viscoelastic-solid like properties.

**Figure 1.**
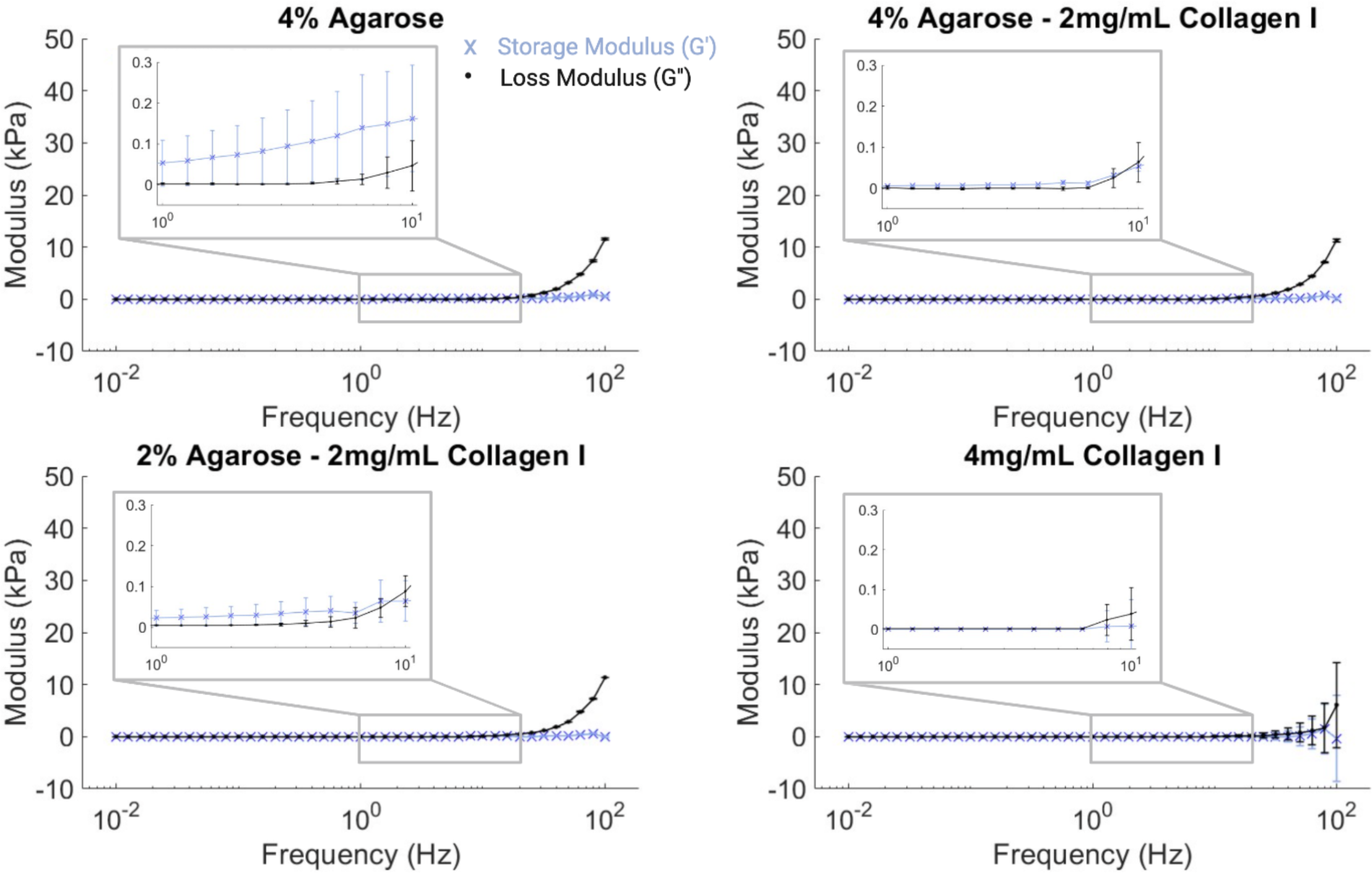
Addition of collagen to agarose reduced the frequency at which the storage and loss moduli cross. The linear equilibrium modulus is defined at the plateau region of *G*′, which occurs at low frequencies for all hydrogels. *G*′ and *G*′′ are shown between 0.01 and 100 Hz and zoomed into the region between 1 and 10 Hz for 4% agarose, 4% agarose – 2 mg/mL collagen, 2% agarose – 2 mg/mL collagen, and 4 mg/mL collagen hydrogels. 3 replicates presented as mean ± SE. Graphs were plotted using Matlab and assembled using Biorender.

All hydrogels displayed a plateau in moduli by 30% strain during strain sweeps (Figure 2). The agarose-only hydrogel required the least strain (1.34±0.32% strain) to achieve the gel-sol transition. Composite 4% agarose – 2 mg/mL collagen and 2% agarose – 2mg/mL collagen hydrogels achieve this transition point at a higher but insignificantly different strain (4.82±1.39% and 4.72±0.74% strain, respectively). The collagen-only hydrogel on the other hand is much more fluid-like and does not reach the gel-sol transition point until 26.8±9.53% strain (*p*<0.05).

**Figure 2.**
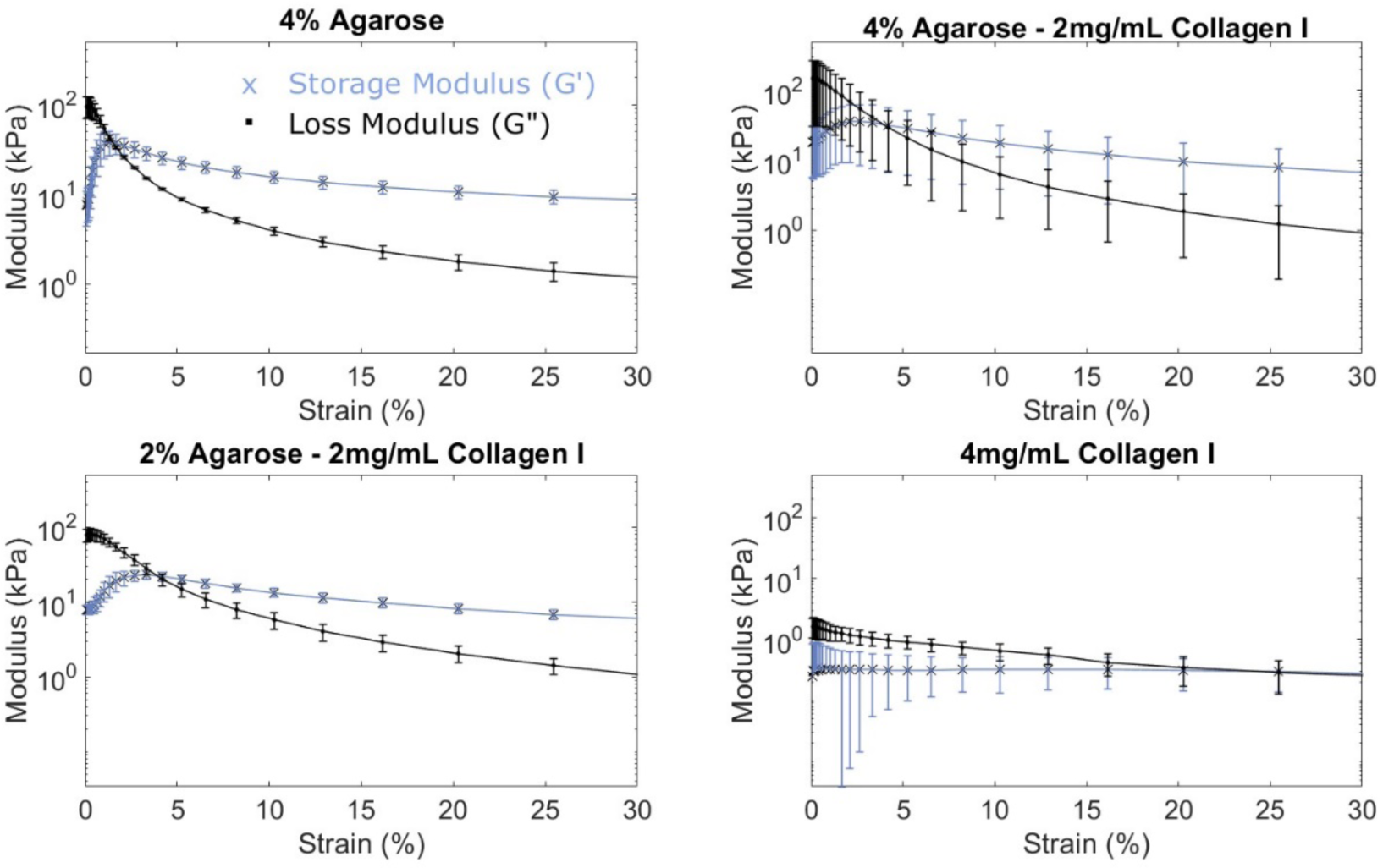
Agarose-only and composite agarose-collagen hydrogels achieved the gel-sol transition point at similar strains. The linear viscoelastic region can be determined with respect to strain where *G*′ and *G*′′ are calculated between 0.1 and 100% strain for all hydrogels. 3 replicates presented as mean ± SE. Graphs were plotted using Matlab and assembled using Biorender.

Within the first 30 minutes of rheometric time sweeps, all hydrogels reached a plateau, indicating that the hydrogels should maintain their mechanical behavior for sustained culture (Figure 3). A storage modulus greater than the loss modulus indicates solid behavior, of which all agarose-containing hydrogels clearly demonstrate at 30 minutes (*p*<0.001). Collagen-only hydrogels, on the other hand, were an order of magnitude softer and demonstrated the opposite effect, though having insignificant differences between storage and loss moduli, in general acting more compliant and fluid-like as compared to agarose-containing hydrogels.

**Figure 3.**
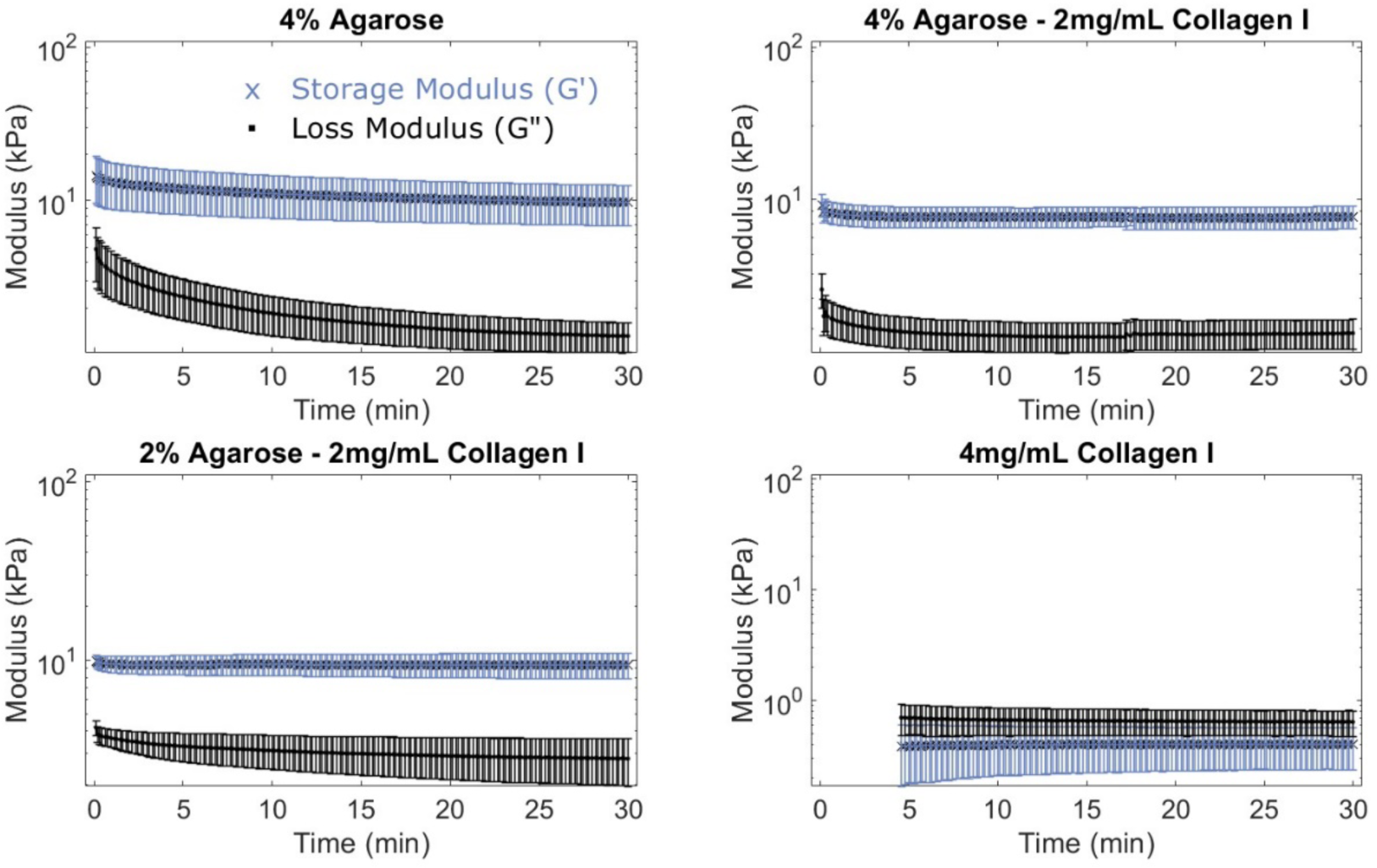
All hydrogel formulations demonstrated stabilized mechanical behavior within thirty minutes. Gel formation was monitored over a period of 30 minutes for all hydrogels. *G*′ and *G*′′ are presented for all formulations; however, the moduli of 4 mg/mL collagen hydrogels did not exceed the limits of instrument sensitivity until after five minutes of gel formation. 3 replicates presented as mean ± SE. Graphs were plotted using Matlab and assembled using Biorender.

In acellular hydrogels, equilibrium modulus was the main interest, as this metric best represents the stiffness of the hydrogels in a swollen state, the same state these samples will undergo during later cell-laden experiments. Stress relaxation under unconfined compression (Figure 4) demonstrated the 4% agarose hydrogels had a similar equilibrium modulus (24.24±5.83 kPa) to the 4% agarose – 2 mg/mL collagen hydrogels (19.72±4.68 kPa). 2% agarose – 2 mg/mL collagen display significantly lower equilibrium moduli (7.86±0.67 kPa) compared to the 4% agarose containing hydrogels (*p*<0.05) in addition to the 4mg/mL collagen hydrogels (1.93±0.46 kPa) showing significantly lower levels than both 4% agarose – 2 mg/mL collagen (*p*<0.01) and 4% agarose (*p*<0.001). Results demonstrate similarities with shear rheometry outputs, as both storage moduli discussed above, and equilibrium moduli here represent the elastic behavior of viscoelastic materials – of which are more prominent in the 4% agarose-containing hydrogels.

**Figure 4.**
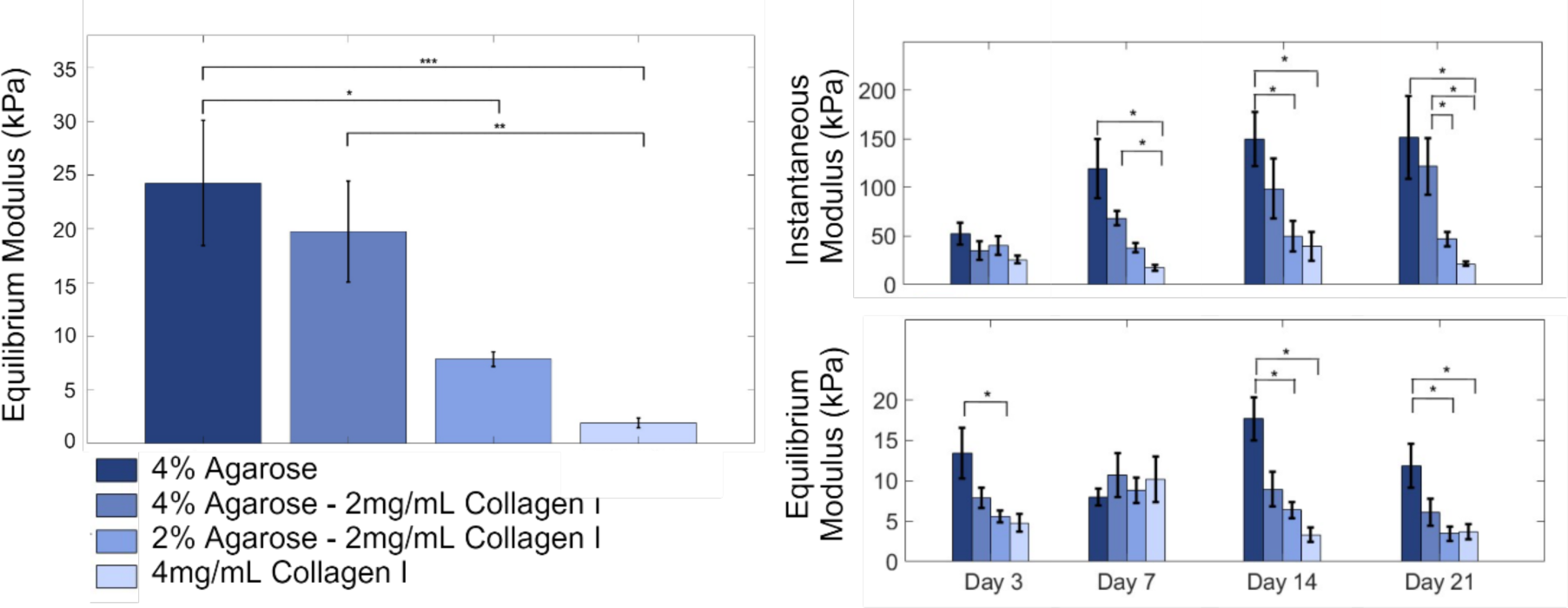
Agarose-only and 4% agarose – 2mg/mL collagen hydrogels displayed similar moduli. Hydrogels (n=6 for 24-hour acellular studies and n=4 for 21-day cell-laden studies) were tested under unconfined, uniaxial compression-based stress relaxation in a randomized order, compared using a one-way ANOVA with post-hoc Bonferroni correction (α=0.05) and presented as mean ± SE. Graphs were plotted using Matlab and assembled using Biorender.

Cell-laden, agarose-containing hydrogels displayed minimal compaction over time, displaying an average diameter increase of 0.09 mm and height decrease of 0.77 mm from initial dimensions. In comparison, collagen-only hydrogels displayed significant changes (*p*<0.05) in height and diameter (0.47±0.26 mm and 0.80±0.32 mm, respectively) after 21 days of chondrocyte culture.

Initially stiffer hydrogels, with respect to the initial equilibrium moduli previously discussed, demonstrate a greater change in instantaneous modulus as compared to softer, collagen dominant hydrogels. Except for these collagen dominant hydrogels, the instantaneous moduli continually increase over 21 days in culture. The final (day 21) instantaneous modulus for 4% agarose hydrogels is 151.77±42.58 kPa, for 4% agarose – 2mg/mL collagen is 121.77±28.93 kPa, for 2% agarose – 2mg/mL collagen is 47.10±7.32 kPa, and for 4mg/mL collagen is 21.63±2.21 kPa (Figure 4).

A similar generalization can be made for the equilibrium moduli over time, though there is much higher variability across time points assessed. Statistically, however, temporal effects were not significant across the 21 days assessed. The introduction and culture of chondrocyte into the hydrogel does cause a decrease in measured equilibrium moduli levels when compared to the acellular hydrogels after 24 hours. In fact, after 21 days in culture, the equilibrium moduli all decrease to roughly half of the initial acellular levels which may be attributed to cellular remodeling of the matrix. The final (day 21) equilibrium modulus for 4% agarose hydrogels is 11.87±2.68 kPa, for 4% agarose – 2mg/mL collagen is 6.11±1.70 kPa, for 2% agarose – 2mg/mL collagen is 3.45±0.91 kPa, and for 4mg/mL collagen is 3.67±0.94 kPa (Figure 4).

### Higher Agarose Concentrations are Associated with Larger Pore Diameters

Hydrogel pore diameters were compared at 100× magnification via SEM (Figure 5). The largest pore diameters were found in 4% agarose hydrogels (19.57±7.61 μm) followed by 2% agarose hydrogels (19.044±7.33 μm), 4% agarose – 2mg/mL collagen and 2% agarose – 2mg/mL collagen pore diameters (16.90±5.90 μm and 16.21±6.55 μm, respectively). Significantly smaller (*p*<0.05) pore diameters were observed in 4- and 2mg/mL collagen samples (9.10±2.68 μm and 12.48±4.10 μm, respectively). Agarose-only hydrogels demonstrated a positive correlation between concentration levels and resultant pore diameters whereas collagen-only hydrogels demonstrated a negative correlation of pore and fiber diameter at 100× and 10,000× magnification levels, respectively (Figure S1).

**Figure 5.**
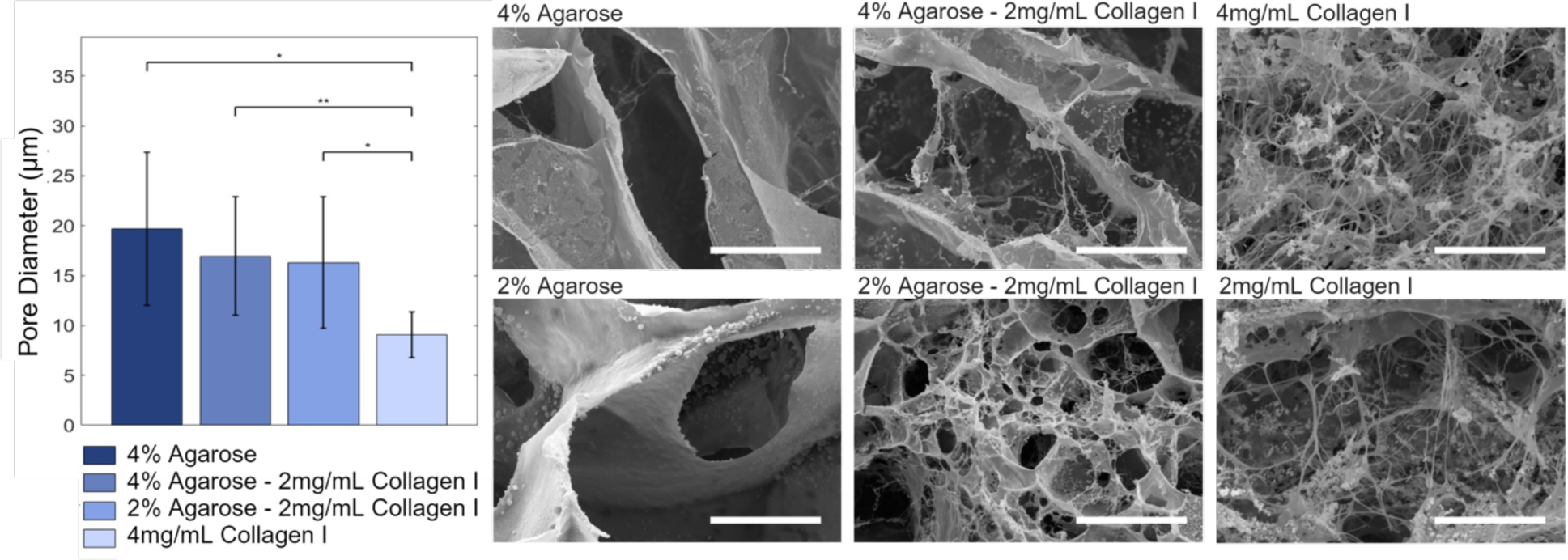
Addition of collagen to agarose reduced pore diameters in SEM images. Pore size was quantified using 3 biological replicate 100× images of lyophilized hydrogels and assessed using a one-way ANOVA with post-hoc Bonferroni correction (α=0.05), presented as mean ± SE. 2000× representative images of hydrogels demonstrate fiber presence and morphology. Scale bars = 20 μm. The graph was plotted using Matlab and assembled using Biorender.

### Composite Hydrogels Display Intermediate Cell Responses between Agarose- or Collagen- only Hydrogels

Since cells were briefly subjected to mechanical and thermal stress during hydrogel formulation methods, cell viability was initially measured directly following cell seeding to establish a baseline. All hydrogels maintained the seeded chondrocytes within their 3-dimensional matrix and demonstrated continued growth, as assessed via resazurin assays and visual inspection under brightfield microscopy. Agarose-only hydrogels demonstrated the quickest initial increase in growth, collagen-only hydrogels initially lagged but eventually demonstrated the largest growth rates (Figure 6). Composite hydrogels meanwhile displayed similar rates and followed the progression of expected cell growth curves through the lag, log, and stationary phases (Figure 6).

**Figure 6.**
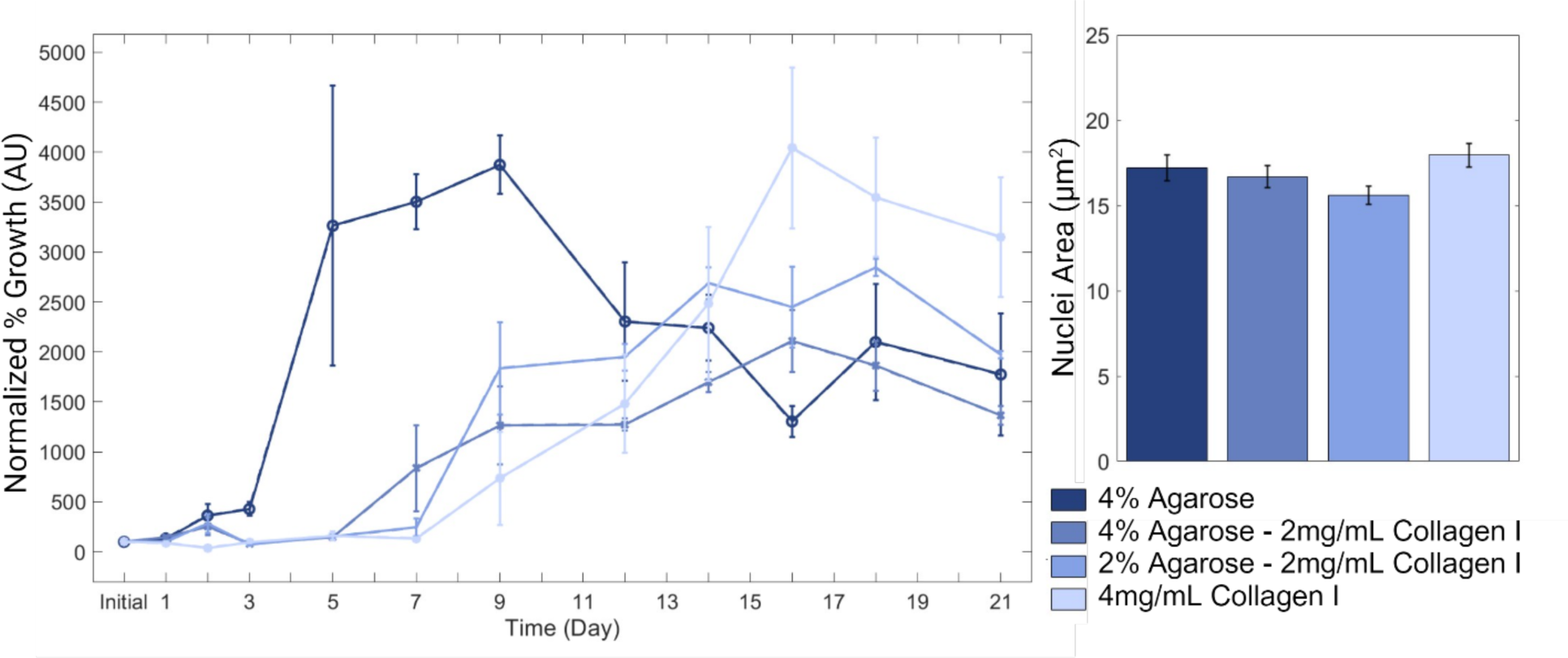
Chondrocyte metabolic activity and nuclear areas, indicators of cell viability and proliferation, demonstrated cell growth across all hydrogels. Cell viability is presented as a percentage of baseline (100%) levels, using 3 biological replicates. Nuclei areas were compared across sections (n=8) from hydrogels (n=2) using a one-way ANOVA with Bonferroni correction (a=0.05) and presented as mean ± SE. Graphs were plotted using Matlab and assembled using Biorender.

The average chondrocyte nucleus area was 16.86±0.32 µm^2^ across hydrogel formulations (Figure 6). Collagen-only hydrogels had the largest areas (17.95±8.24 µm^2^) with agarose following (17.22±10.45 µm^2^). Interestingly, as agarose concentration decreased and collagen content increased in the composite hydrogels, so did nuclear areas (16.68±8.25 µm^2^ for 4% agarose – 2mg/mL collagen hydrogels and 15.60±9.33 µm^2^ for 2% agarose – 2mg/mL collagen hydrogels).

### Hydrogel Formulation is Not a Significant Driver in Observed sGAG Content

sGAG was analyzed in terms of absolute content. Interestingly, both composite hydrogels demonstrated a brief decrease in sGAG content between days 3 and 7 before continuing to show continuous increases in content. DMMB results demonstrate on day 3, sGAG content in 4% agarose – 2mg/mL collagen hydrogels were statistically higher than other groups (Figure 7). Other than this initial case, however, the temporal influence was the main driver on sGAG content (*p*<0.01) while formulation alone was not significant driver (*p*=0.7), indicating that the addition of agarose in the composite hydrogels does not hinder the ability of chondrocytes to secrete sGAG in vitro at the levels tested.

**Figure 7.**
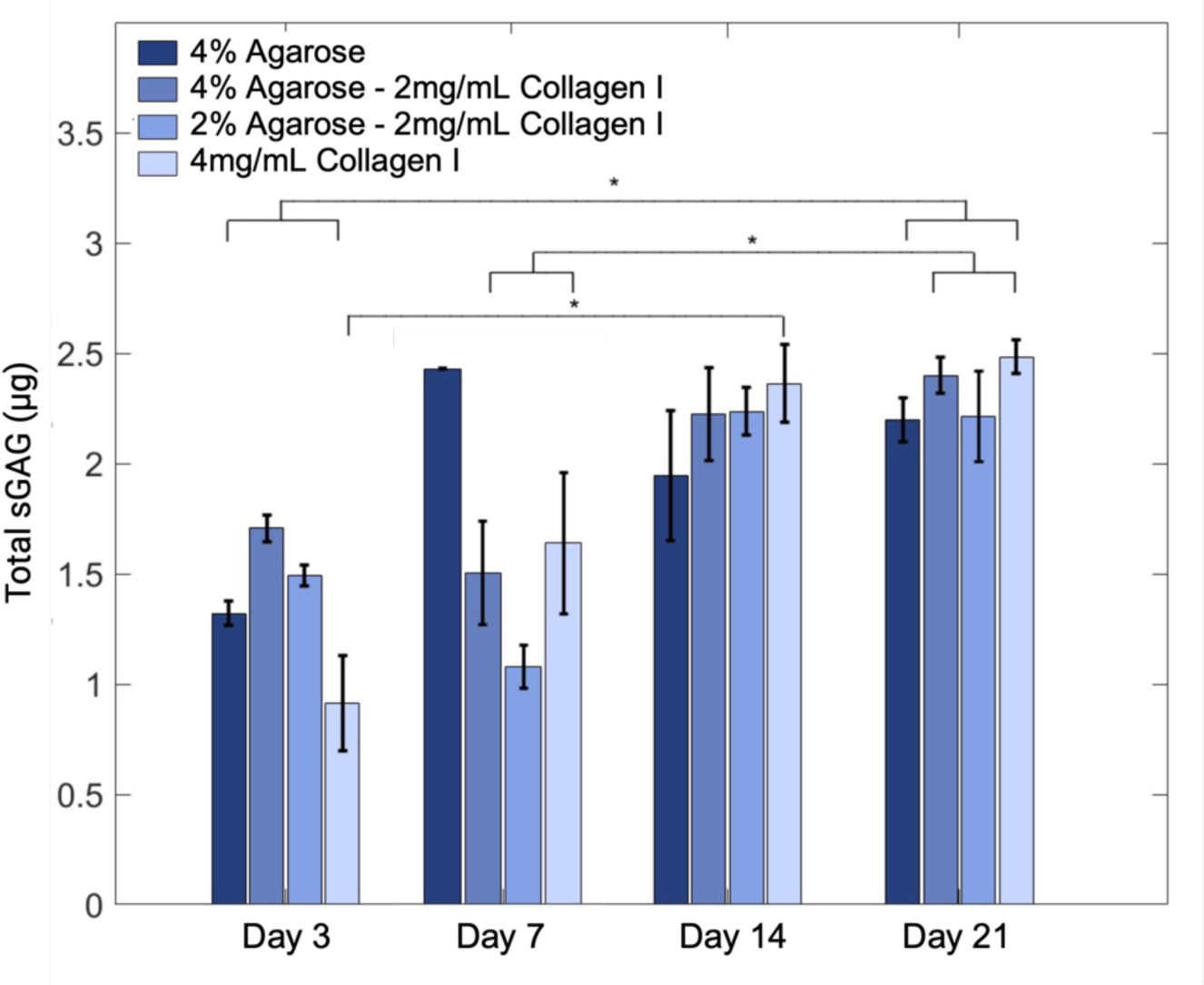
Collagen-containing hydrogels demonstrated an increase in sGAG content that plateaued by day 21. One-way ANOVA with post-hoc Bonferroni corrections (α=0.05) were used to detect temporal and hydrogel formulation influences on sGAG content in hydrogels (n=3) and results are presented as mean ± SE. Graphs were plotted using Matlab and assembled using Biorender.

## DISCUSSION

Biomaterials meeting the mechanical and biochemical requirements for functional environments during cell culture are necessary for mechanotransduction. This study aimed to evaluate agarose-collagen composite hydrogels as a simple, effective option for chondrocyte mechanobiology studies. Four hydrogel formulations were assessed, including a high-concentration, 4% agarose hydrogel, two common-level collagen concentrations with varied agarose (4% agarose – 2mg/mL collagen and 2% agarose – 2mg/mL collagen), and a 4mg/mL collagen hydrogel. Both acellular and prolonged cell-laden studies indicate that high-concentration agarose blended with collagen hydrogels may be a suitable environment for extended, 3-dimensional chondrocyte culture.

### Mechanical Properties describe the Hydrogel Structure-Function Relationship

Of the sweeps performed, all acellular hydrogels demonstrated linear viscoelastic characteristics that fall within the bounds of common parameters used in loading studies ^34,40^, a critical factor when developing a hydrogel material meant for culture in physiologically relevant, dynamic conditions. The addition of collagen into agarose did not significantly change the rheologic properties of the hydrogel though it did slightly reduce the elasticity. The blending of collagen and agarose at these concentrations therefore do not seem to impair bond formation during gelation, though it is possible that at higher collagen concentrations, fiber aggregation might interfere with gelation, leading to macroscopic defects, as suggested by the results found by Cambria, *et. al.* ^28^ and SEM images of collagen-only samples.

Compression testing further described sample mechanical characteristics. Acellular 4% agarose hydrogels were the stiffest material after gelation, followed by 4% agarose – 2 mg/mL collagen, which both fall within the lower bounds of native cartilage PCM ^19,20^. The moduli values of the composite hydrogels demonstrate a dependence on the volume fractions of the agarose and collagen, in which hydrogels with higher agarose content showed improved mechanical properties, reflecting changes to hydrogel moduli reported in other studies. 0.5% agarose hydrogels have an equilibrium modulus ranging from 5 to 8 kPa ^29^, 2% agarose hydrogels at 14 kPa ^41^, 3% agarose hydrogels at roughly 20 kPa ^14^, and 4% agarose hydrogels at 40 kPa ^41^. Meanwhile, collagen-only hydrogels demonstrated substantially lower equilibrium moduli following compression ^42^, like that of 2.3 mg/mL collagen hydrogels, which have a modulus of only 0.1 kPa ^43^.

Cell-laden agarose-containing hydrogels presented the most structural stability over time, in part due to the lack of agarose production by chondrocytes. Since the cells do not produce any enzymes that break down agarose, the material does not erode quickly ^30^. Collagen-based hydrogels, on the other hand, are well known to contract over time, decreasing upwards of 70% in diameter ^23,27^. Supplementing observations from rheologic testing, the low levels of contraction observed in the composite formulations demonstrated how the agarose component dominated over collagen to maintain the hydrogel shape with time. This presents an advantage for studies in which hydrogels need to be cultured for long durations or undergo loading to mimic physiologic conditions. advantage for studies in which hydrogels need to be cultured for long durations or undergo loading to mimic physiologic conditions.

In general, the relaxation behavior of the hydrogels seems to be dominated by the agarose composition. The observed increase in moduli over time may be due to greater fiber engagement or increased matrix deposition as the cells continue to proliferate ^32^. While trends observed in this study follow those elsewhere ^14,28,41,43^, discrepancies in absolute values may be due to differences in experimental setup, including plate geometries, preconditioning tare loads, and relaxation times in compression testing.

Pore diameter sizes seem to follow the same trend as equilibrium moduli, in which 4% agarose hydrogels, the material with the largest pore diameter, also demonstrate the highest moduli. This related change in pore geometry and subsequent material property are likely attributable to the reduction of hydrogel bonding with decreased agarose concentrations ^26,29,44,45^. The composite hydrogels demonstrate similar porosities and moduli but are both lower than agarose-only hydrogels. These composites are double networked since they consist of contrasting component properties and molar concentrations. The differences in crosslinking mechanisms between materials within these hydrogels likely influence the observed trends in pore and modulus values ^46–48^. Collagen-only hydrogels, meanwhile, demonstrate a negative relationship between porosity and moduli, possibly mediated in part by bond strengths, as there exists a positive relationship between collagen concentration and ionic strength ^45^ and further a positive relationship between increases in ionic bond strength leads and fiber networks connectivity ^30,49^, leading to smaller pore and fiber diameters. Furthermore, variability between samples can arise and is demonstrated in literature depending on collagen source, polymerization temperature, and pH, all of which affect crosslinking and cellular interactions ^44,50–52^.

### Cell and Extracellular Matrix Responses describe the Hydrogel Biofunctionality

In this study, the stiffest hydrogel, 4% agarose-only, led to the quickest and most dramatic increase in proliferation, as measured through mitochondrial activity, although the 4mg/mL collagen-only hydrogels reached similar growth rates within two weeks of culture. Previous reports are inconclusive towards the relationship between hydrogel viscosity and cell proliferation over a 21-day period. For example, Lee, *et. al.*, found that higher viscosities led to higher proliferation rates ^53^ whereas Cambria, *et. al.*, found the opposite ^28^. Potential explanations for the contradictory results may be the influence of contact inhibition or biochemical signaling differences across biomaterials (gelatin blends versus agarose-collagen blends, respectively). Since mammalian cells will not bind to agarose polysaccharides, it is possible that the embedded cells were able to begin proliferating much sooner in a free-floating state as compared to the collagen-containing hydrogels that allowed the cells to take time and bind to fibers prior to entering the exponential growth phase. The composite hydrogels, on the other hand, both resulted in much steadier and more predictable growth curves, a feature that may be advantageous for mechanobiology studies.

The average nuclei diameter observed resembles dimensions of chondrocyte nuclei reported at earlier time frames of culture ^28,54,55^ and aspect ratios remain circular-like in 2-dimensional images after 21 days in culture (Figure S2), indicating the hydrogel material is not significantly influencing the cells to dedifferentiate and may be suitable in preventing terminal differentiation ^56^. While it is well established that cells are sensitive to substrate stiffnesses in 2-dimensional culture ^57–60^, the influence of substrate stiffness is less understood in 3-dimensional culture ^61^. Additional studies with an increased focus on fluorescence-based microscopy would provide further insight to the morphologies during 3-dimensional culture.

No significant differences in sGAG content by time point were observed among the formulations after day 3. Over the course of the study, all hydrogels continued to demonstrate greater sGAG content, suggesting continual matrix synthesis. A lack of consensus exists regarding sGAG content in composite hydrogels. While some reports show higher collagen content associated with higher sGAG content ^28^, others report the opposite effect, in which a low collagen content led to higher sGAG content ^31^. However, as previously referred to, these differences could also be attributable to discrepancies among collagen sources and associated influences on cell behavior.

## CONCLUSION

This study focused on the structural and biofunctional properties of agarose-collagen composite hydrogels. While this study had a low sample size per group due to time constraints on mechanical testing, material availability, and the living nature of the material, it has allowed us to establish the structural properties and compatibility of 4% agarose – 2mg/mL collagen hydrogels. Future studies including gene and protein expression analyses would provide deeper insight into chondrocyte responses to the physical and mechanical cues presented by the composite hydrogels. These hydrogels can be easily and quickly produced, mechanical shear and compressive properties show promising behavior as stable, long-term environments, microscopy images demonstrate homogenous and interconnected networks for cell growth, and preliminary cell-laden studies demonstrate continual proliferation, matrix deposition, and maintained morphology. Overall, the 4% agarose – 2mg/mL collagen hydrogel formulation showed potential in the context of chondrocyte mechanobiology studies.

## ACKNOWLEDGEMENTS

### Funding

This research was supported by the National Science Foundation (NSF) Award Number 2149946.

### Author Contributions

CMZ: conceptualization, data curation, formal analysis, investigation, methodology, software, validation, visualization, writing - original draft, review, and editing

CBA: data curation, formal analysis, investigation, methodology, software, writing - original draft, review, and editing

AC: data curation, methodology, software, writing – review and editing LS: methodology, resources, writing - review and editing

DDC: conceptualization, funding acquisition, methodology, project administration, resources, supervision, writing - review and editing

### Ethics and Integrity Policy Statements

Data availability statement: Collected data is made available using a Harvard Dataverse repository.

Funding statement: This research was supported by the National Science Foundation (NSF) Award Number 2149946.

Conflict of interest disclosure: The authors have no conflicts of interest to disclose.

Ethics approval statement: To obtain primary murine cells, animals were euthanized following institutional approval (IACUC protocol 2104002138).

Patient consent statement: N/A

Permission to reproduce material from other sources: N/A Clinical trial registration: N/A

## Abbreviations

DAPI 4: 6-diamidino-2-phenylindole
DMEM: Dulbecco’s modified Eagle medium
ECM: extracellular matrix
FBS: fetal bovine serum
FE-SEM: field emission scanning electron microscopy
LVR: linear viscoelastic region
PBS: phosphate buffered saline
PCM: pericellular matrix
sGAG: sulfated glycosaminoglycan

