## Supplementary material for "Characterization of Composite Agarose-Collagen Hydrogels for Chondrocyte Culture": Figure S1

**SUPPLEMENTAL INFORMATION**

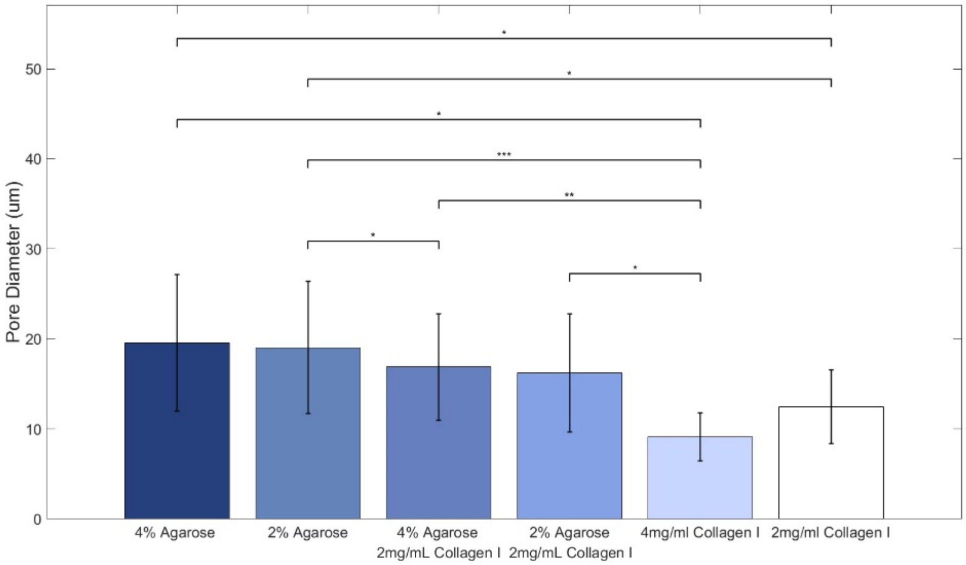


**Figure S1. The dilution of agarose in agarose-only and composite hydrogels demonstrated a positive relationship with measured pore diameters, whereas in collagen-only hydrogels, an inversed relationship exists.** Fiber diameters were quantified using 100× images of lyophilized hydrogels (n=3), assessed using one-way ANOVA with Bonferroni corrections (a=0.05) and presented as mean ± SE. Graphs were plotted using Matlab and assembled using Biorender.
