## Supplementary material for "Characterization of Composite Agarose-Collagen Hydrogels for Chondrocyte Culture": Figure S2

**SUPPLEMENTAL INFORMATION**

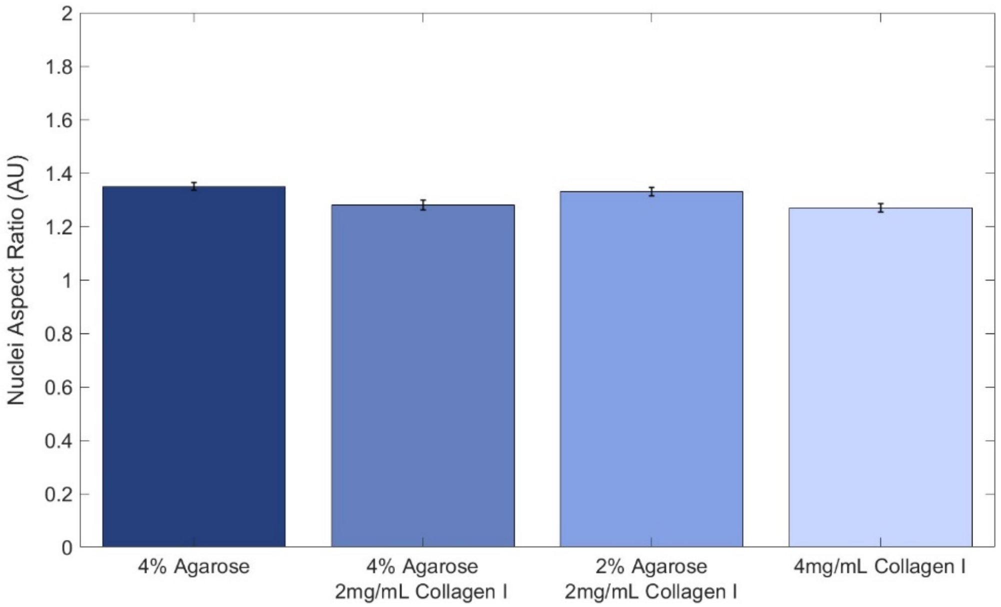


**Figure S2. Chondrocyte nuclei aspect ratios are not impacted due to hydrogel formulation after 21 days of culture.** DAPI-stained hydrogel sections under confocal microscopy provided information on nuclear morphology and spatial orientation. Sections (n=8) were obtained from hydrogels (n=2), assessed using a one-way ANOVA with Bonferroni correction (a=0.05) and presented as mean ± SE. Graphs were plotted using Matlab and assembled using Biorender.
